# Linguistic information compensates for age-related decline in attentional filtering

**DOI:** 10.64898/2026.01.08.698329

**Authors:** Alice Vivien Barchet, Andrea Bruera, Johanna M. Rimmele, Jonas Obleser, Gesa Hartwigsen

## Abstract

As we age, understanding speech in social situations imposes an increasingly difficult challenge to the auditory system. However, the attentional mechanisms underlying age-related speech comprehension difficulties in multitalker situations remain unclear. We collected EEG signals while 63 normal hearing participants from 19 to 71 years performed a speech comprehension task involving a multitalker paradigm at individually adjusted target-to-distractor ratios. Combining trial-resolved multivariate temporal response function modeling with detailed behavioral comprehension responses, we provide a window into lower-level impairments and higher-level compensatory mechanisms across the adult life span. Neuro-behavioral correlations on a trial-by-trial level provide direct evidence for increased distractor representation underlying reduced behavioral performance in late adulthood. This points towards increased distractibility as a potential mechanism underlying age-related speech comprehension deficits. Additionally, at the behavioral and neural levels, we show that older adults relied more on higher-level linguistic information. Finally, we show that an increased reliance on word-level linguistic information may compensate for increased distractor tracking and adaptively support comprehension performance across the adult life span. Specifically, we directly show that increased reliance on higher-level processing can offset age-related impairments in attentional filtering.

## Introduction

In everyday life, spoken speech is often masked by noise or competing speech streams in the surrounding, creating challenging listening situations, particularly for older adults^1–3^. However, it remains unclear why older adults’ speech comprehension is impaired in multitalker contexts. Disentangling the causes for impaired speech comprehension is particularly important as impaired hearing abilities are associated with poor cognitive and psychosocial functioning in late adulthood^4,5^.

Healthy aging results in many changes in sensory processing, including hearing loss and reduced auditory temporal resolution^6–10^. At the same time, speech comprehension remains largely preserved across the adult life span^11,12^. However, older adults often experience difficulties in perceptually demanding listening conditions, such as competing speech situations^11–13^. Successful speech comprehension in challenging listening situations requires separating the to-be-attended target stream from one or more to-be-ignored distractor streams. Speech comprehension seems to be related to low-frequency cortical activity that temporally aligns with the speech signals^14–16^. In competing speech comprehension, neural tracking of acoustic properties of the to-be-attended signal has been shown to be selectively enhanced^17–21^. Recent evidence suggested that this attentional filtering is particularly susceptible to senescent changes, potentially due to impaired neural inhibition^22,23^.

Behavioral studies have suggested that older adults rely more on contextual cues than younger adults to maintain speech comprehension ability despite peripheral declines^13,24,25^. Correspondingly, processes involved in the activation of contextual knowledge have been shown to be relatively unaffected by aging^26,27^. Yet, neural results employing event-related potentials have revealed weaker responses to semantic context in older adults than in younger adults^28–30^. Interestingly, recent methodological advances have enabled researchers to investigate neural speech tracking in natural speech on several hierarchical levels, including acoustic, phonemic, and semantic information in a unified encoding framework^31–34^. Here, neural results concerning the representation of semantic information across the adult life span have revealed heterogeneous results, suggesting that semantic information might be represented more strongly^35^, similarly^36^, or less strongly^37^ in older adults’ brain signals. In contrast, the neural tracking of acoustic properties is typically observed to be stronger in older adults^38–40,37^. However, it is unclear whether this increased tracking of acoustic properties serves a compensatory purpose or if it reflects increased processing noise^22,38,41–43^.

To disentangle neural mechanisms of age-related speech comprehension deficits from compensatory mechanisms, it is crucial to link measures of neural tracking with behavioral comprehension performance. However, such a link is missing in most previous studies. Most studies investigating speech tracking in naturalistic contexts lack detailed behavioral responses and are therefore unable to relate neural tracking to time-resolved variations in comprehension success^34–37,39^. Here, we combined time-resolved behavioral outcome measures and EEG using a naturalistic competing speech paradigm to probe the behavioral relevance of acoustic and higher-level processing across the adult life span. In contrast to previous studies that used natural listening paradigms, we asked participants to listen to short sentences and repeat the words they understood. This allowed the analysis of comprehension dynamics using a fine-grained, word-by-word intelligibility measure. Additionally, task difficulty was individually adjusted based on the individual speech reception threshold (SRT) as measured by an adaptive staircase procedure to control for effects of peripheral constraints. As a result, the average comprehension performance was balanced across participants to allow for conclusions regarding the use of different strategies to achieve similar performance levels. Using multivariate temporal response function (TRF) modeling, we provide insight into the trial-wise neural tracking strength of acoustic and linguistic speech representations for the target and distractor streams, as well as their relationship to behavioral comprehension performance and changes of this relationship across the adult life span.

By employing trial-resolved analyses on the behavioral and neural levels, we addressed the overarching research question of the mechanisms underlying age-related changes in speech comprehension in multitalker environments. Investigating speech features at different hierarchically organized levels allowed us to disentangle how lower-level and higher-level linguistic mechanisms support comprehension performance in a challenging and naturalistic multitalker paradigm. Here, lower-level mechanisms refer to phonetic (i.e., envelope) and phonological (i.e., word and phoneme onsets) processing. Higher-level mechanisms refer to responses to probabilistic features, such as word frequency, word surprisal, and word entropy.

Based on previous evidence suggesting age-related changes in the temporal resolution and hearing acuity, we expected older adults to display lower-level impairments potentially affecting their attentional filtering strategies. Additionally, we hypothesized that in a multitalker environment like the one we tested here, older adults would engage in a compensatory recruitment of higher level mechanisms in speech comprehension. Specifically, older adults should demonstrate a stronger reliance on higher-level information on the behavioral and the neural levels.

Our results demonstrate that the neural tracking of the distractor stream may serve as a behaviorally relevant indicator of age-related impairments in stream segregation. At the same time, behavioral and neural results indicate a stronger reliance on higher-level target processing with increasing age. Collectively, our results highlight the interplay of lower-level impairments and higher-level compensation across the adult life span.

## Results

### Behavioral Results

#### Both lower-level and higher-level features predict comprehension performance

In a generalized linear mixed model, we predicted word-by-word comprehension performance from word-level audibility, surprisal, and entropy for the target stream. Results revealed that audibility, surprisal, as well as their interaction, predicted word repetition performance. Figure 1 B displays the interaction between word audibility and surprisal, indicating that the negative effect of surprisal is stronger at low levels of audibility, with higher comprehension for low surprisal (i.e., more predictable) words. Word entropy positively predicted comprehension performance. While this effect seems counterintuitive, it is consistent with previous studies showing more efficient lexical processing in contexts of high uncertainty^44,45^. This effect could indicate that the allocation of attention as well as the pre-activation of semantic features is increased in contexts of high uncertainty. Overall, there were negative effects of age and PTA on comprehension performance. Complete results for the generalized linear mixed model can be retrieved from Table 1.

**Figure 1:**
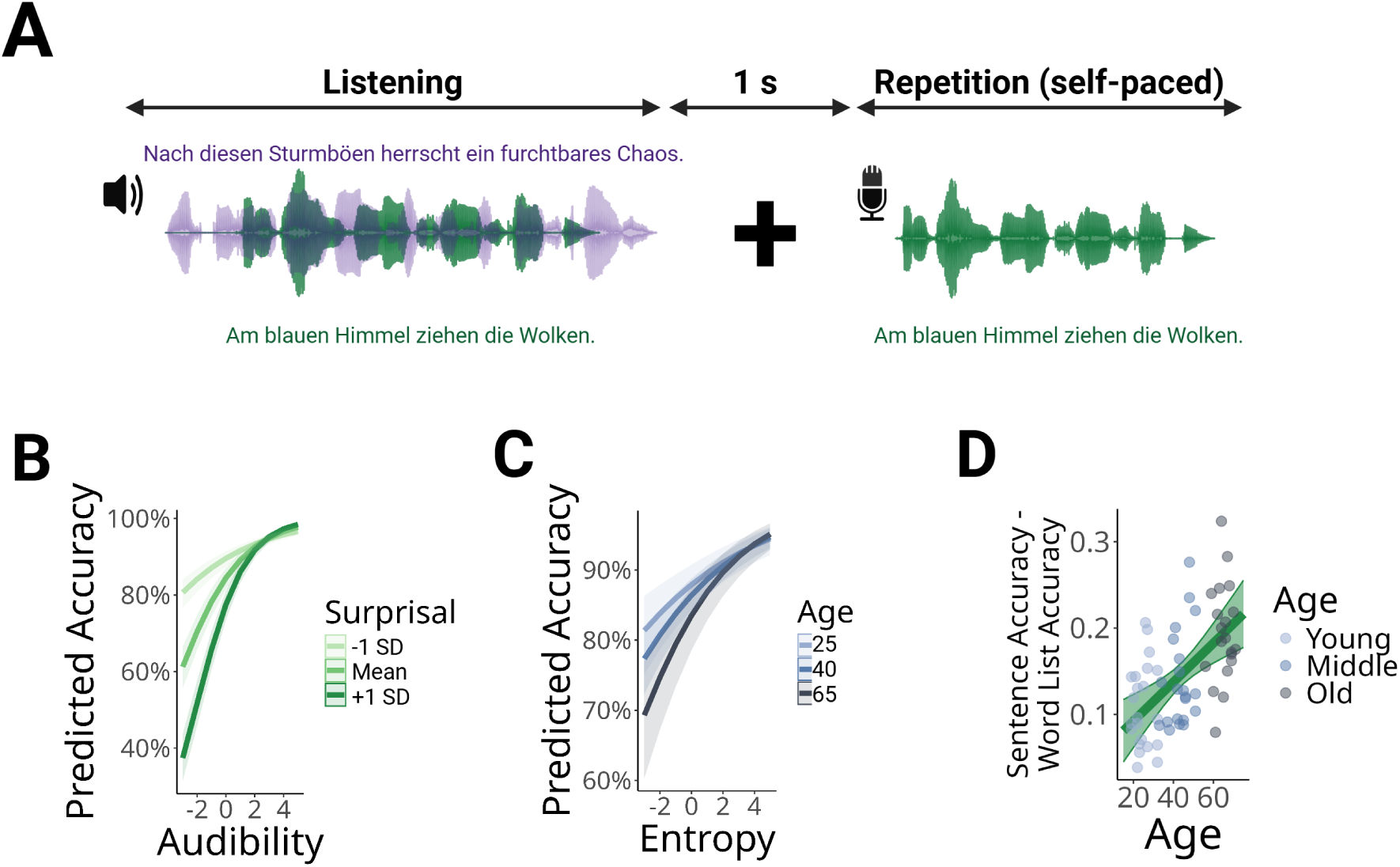
Behavioral analysis. A: Behavioral Paradigm. Participants listened to 240 trials comprised of one target and one distractor sentence. They subsequently repeated the target sentence. English translations of the sentences: After these squalls, there is a terrible chaos (purple), Clouds gather in the blue sky (green). B: Effects of surprisal and audibility on comprehension accuracy. Surprisal and audibility jointly predict comprehension accuracy, with a bigger effect of surprisal at low levels of audibility. C: Interaction between age and word entropy. Word entropy had a stronger positive effect on comprehension accuracy in older adults (>40 years of age) than in younger adults (<40 years of age). D: Comprehension gain from sentence context and age. The accuracy difference between the main experiment and an additional block of word lists was larger with increasing age. Age was used as a continuous predictor in all analyses and age groups were used for illustrative purposes only.

**Table 1:** Model outputs predicting comprehension performance.

|  | **Estimate** | **Std. Error** | **CI** | **z** | **p** |
| --- | --- | --- | --- | --- | --- |
| **a) Behavioral effects** |  |  |  |  |  |
| Age | -0.53 | 0.08 | [-0.7, -0.37] | -6.43 | <.001 |
| Audibility | 0.41 | 0.01 | [0.38, 0.44] | 27.77 | <.001** |
| Surprisal | -0.46 | 0.02 | [-0.5, -0.42] | -22.29 | <.001** |
| Audibility:Surprisal | 0.17 | 0.02 | [0.14, 0.2] | 11.07 | <.001** |
| Word frequency | 0.15 | 0.02 | [0.1, 0.19] | 6.01 | <.001** |
| Word entropy | 0.21 | 0.02 | [0.17, 0.24] | 10.95 | <.001** |
| Audibility:Age | 0 | 0.01 | [-0.03, 0.02] | -0.39 | .756 |
| Surprisal:Age | 0 | 0.02 | [-0.03, 0.03] | 0.16 | .905 |
| Entropy:Age | 0.04 | 0.02 | [0.01, 0.07] | 2.8 | .009** |
| **b) Lower-level TRF Fits** |  |  |  |  |  |
| Acoustic target | 0.01 | 0.01 | [-0.01, 0.03] | 0.89 | .478 |
| Acoustic distractor | 0.03 | 0.01 | [0.01, 0.05] | 2.45 | .022* |
| Onsets target | 0.04 | 0.01 | [0.02, 0.07] | 3.35 | .002** |
| Onsets distractor | -0.01 | 0.01 | [-0.03, 0.01] | -0.86 | .478 |
| Acoustic target:Age | 0 | 0.01 | [-0.03, 0.02] | -0.08 | .933 |
| Acoustic distractor:Age | 0.01 | 0.01 | [-0.02, 0.03] | 0.46 | .730 |
| Onsets target:Age | -0.01 | 0.01 | [-0.04, 0.01] | -1.13 | .370 |
| Onsets distractor:Age | -0.03 | 0.01 | [-0.06, -0.01] | -2.85 | .008** |
| **c) Higher-level TRF Fits** |  |  |  |  |  |
| Word target | 0.03 | 0.01 | [0.01, 0.06] | 2.7 | .011* |
| Word target:Age | 0 | 0.01 | [-0.02, 0.03] | 0.53 | .698 |
| **d) Control Variables** |  |  |  |  |  |
| Pure tone average | -0.07 | 0.02 | [-0.1, -0.04] | -4.25 | <.001** |
| Target-to-distractor ratio | 0.49 | 0.01 | [0.47, 0.51] | 55.89 | <.001** |
| Word length | 0.08 | 0.01 | [0.07, 0.1] | 11.64 | <.001** |
| Word order | -0.13 | 0.01 | [-0.15, -0.11] | -13.35 | <.001** |
| Sentence length | -0.1 | 0.11 | [-0.32, 0.12] | -0.9 | .478 |
| Control accuracy | 0.24 | 0.07 | [0.1, 0.38] | 3.26 | .002** |
| Speech reception threshold | 0.29 | 0.03 | [0.23, 0.35] | 9.11 | <.001** |

\*\* p < .01, \* p < .05 (FDR corrected), N = 63

#### Stronger behavioral reliance on entropy in older adults

Additionally, the model revealed an interaction between age and word entropy. This interaction indicates that the effect of word entropy on comprehension was stronger with increasing age (Figure 1 C). More precisely, there seems to be a stronger negative effect of low word entropy on comprehension performance with increasing age. This effect suggests that older adults’ comprehension performance is modulated more strongly by the linguistic context.

#### Comprehension gain from sentence context increases with age

To quantify how much each participant’s behavioral accuracy benefited from sentence context, we calculated standardized differences between the mean accuracy in the main experiment and an additional block of lists of unrelated words. A multiple linear regression model predicting these difference scores from age revealed a positive correlation between age and the accuracy gained from sentence context (Estimate = 0.57, SE = 0.15, t = 3.71, p < .001). The effect is visualized in Figure 1 D. Thus, in a noisy environment, older adults seem to benefit more than younger adults from the richer linguistic context provided by meaningful, coherent sentences as opposed to unrelated sets of words.

### Lower-level TRF model fits

#### Significant lower-level tracking for the target and distractor streams

To investigate the neural tracking of lower-level features, we conducted a TRF analysis predicting the EEG signals from acoustic features and word and phoneme onsets. Acoustic features and onsets were shuffled separately to disentangle acoustic processing and word and phoneme segmentation. As displayed in Figure 2 A, the lower-level features were significantly tracked for the target and distractor streams. Accordingly, significant encoding model performance with increased model fit compared to a shuffled null distribution was observed for the acoustic features and the word and phoneme onsets. The topographies reveal the strongest encoding accuracies in frontal and central sensors.

**Figure 2:**
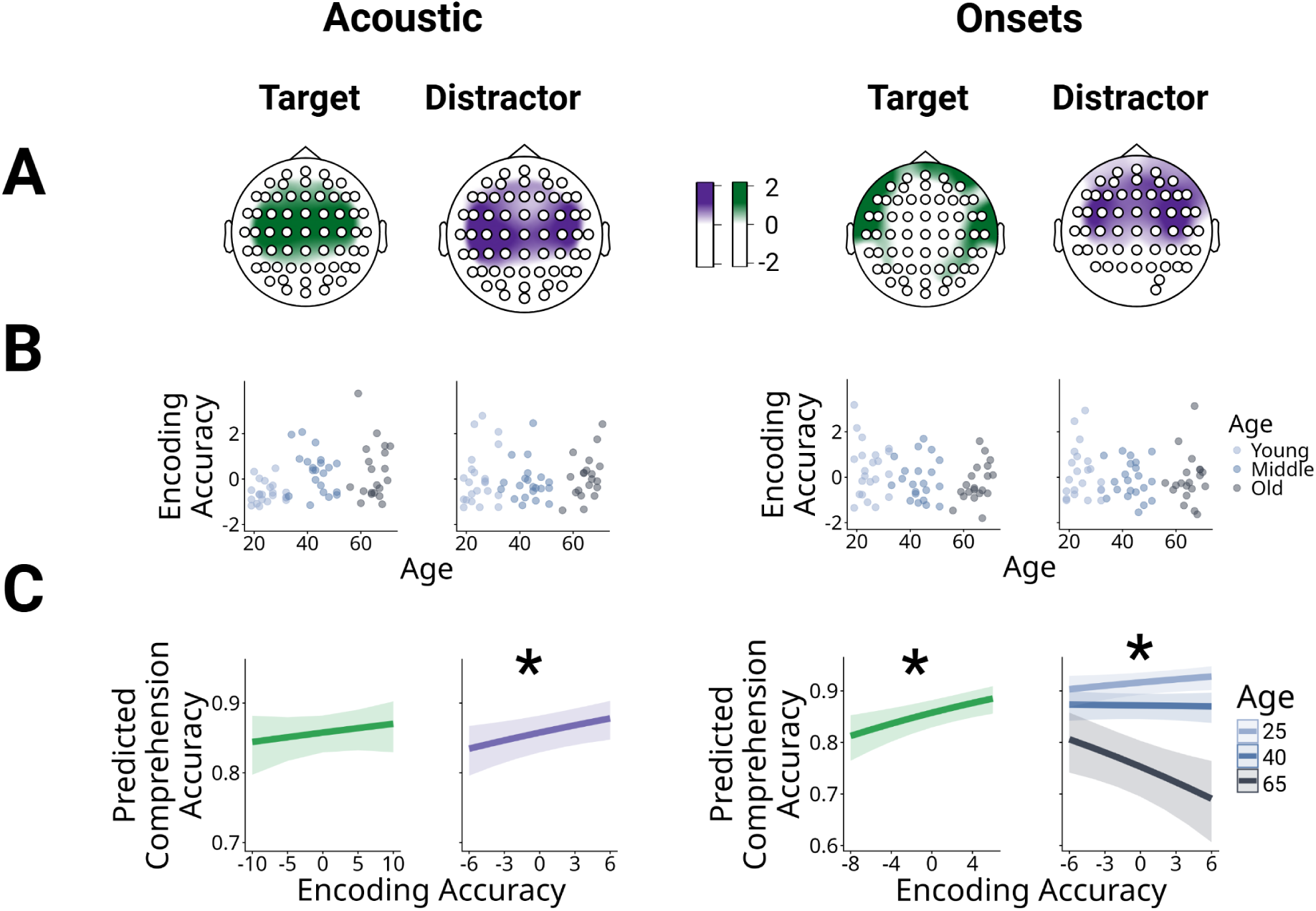
Neural tracking and neuro-behavioral correlations for the lower-level features. A: Topographies for the encoding accuracies. Significant electrodes in mass-univariate t-tests are marked by white dots. B: Age effects on encoding accuracy. No significant main effects of age on encoding accuracy strength were observed. C: Neuro-behavioral correlations and age interactions. Linear mixed model predictions of the effects of trial-wise neural encoding accuracy on comprehension accuracy. Distractor acoustic features and target onsets showed a positive relationship with comprehension accuracy. A significant interaction between the encoding model fits and age emerged for the distractor onsets, suggesting that older adults’ comprehension performance is more negatively influenced by the neural tracking of distractor features.

#### No effects of age on lower-level encoding accuracy

To assess the effects of age on encoding accuracy, we used the mean encoding accuracy of the lower-level models across all sensors. We used multiple linear regression models predicting the mean encoding accuracy for every participant from age. The analysis was controlled for the participant-specific speech reception threshold (SRT), the accuracy in the main experiment and in the control condition, and hearing ability as measured by the pure tone averge (PTA). The control condition consisted of well intelligible sentences presented at SRT +10 dB. There were no significant associations between age and the strength of lower-level encoding accuracies.

#### Stronger lower-level encoding is related to better comprehension

Using a generalized linear mixed model, we predicted word-by-word comprehension accuracy from the trial-wise TRF model fits, as well as their interactions with age. We observed positive effects of lower-level tracking on comprehension performance for the target and distractor streams. More precisely, the neural tracking of target word and phoneme onsets and the tracking of distractor acoustics were positively related to comprehension accuracy. The model predictions are visualized in Figure 2 C. First, these effects show that word and phoneme segmentation of the target stream is crucial for comprehension. Second, increased acoustic tracking of the distractor stream was related to improved comprehension performance across the full age range. This could indicate that an initial acoustic analysis of the distractor stream and segmentation of the target stream are beneficial for stream segregation and comprehension performance.

#### Increased age correlates with stronger distraction

The generalized linear mixed model revealed a significant interaction between the effects of age and the neural tracking of word and phoneme onsets in the distractor stream on comprehension performance. A post-hoc simple slopes analysis revealed a significant negative effect of the neural tracking of distractor onsets in older adults (slope of distractor onsets when age = 65: Estimate = −0.05, SE = 0.02, z = −2.65, p = .01*), but not in younger (slope of distractor onsets when age = 25: Estimate = 0.03, SE = 0.02, z = 1.60, p = .11) or middle aged adults (slope when age = 40: Estimate = −0.00, SE = 0.01, z = −0.22, p = .82).

Thus, a stronger neural representation of distractor onsets had a negative effect in older adults, but not in younger adults. On this intermediate level of (sub-)lexical segmentation, older adults’ comprehension performance seems to be negatively affected by the neural tracking of the distractor sentence. This indicates that an increased allocation of attention to the distractor stream occurring at an intermediate level might underlie older adults’ speech comprehension difficulties. The effect is visualized in Figure 2 C. Although the tracking of distractor acoustics was positively related to comprehension, we observed a negative effect of distractor onsets in older adults. This might indicate that, while acoustic analysis is required for comprehension, tracking of the intermediate onset level marks a distraction effect that is detrimental for comprehension.

### Higher-level TRF model fits

#### Word-level features are significantly tracked for the target stream

To investigate the neural tracking of higher-level features, we conducted a separate TRF analysis predicting the residualized EEG responses from word-level (word frequency, surprisal, and entropy) and phoneme-level (phoneme surprisal and phoneme entropy) probabilistic features. As displayed in Figure 3 A, the word level probabilistic features were significantly tracked for the target stream. This was expected based on previous results and indicates a successful allocation of attention towards the target stream, at least in the majority of trials (see [46] for an exploration of the neural tracking of the distractor stream resolved by comprehension accuracy). The phoneme level probabilistic features were not significantly tracked. Similarly, the higher-level features were not significantly tracked for the distractor stream. Therefore, we did not include the distractor higher-level tracking and the phoneme level tracking in the subsequent analyses.

**Figure 3:**
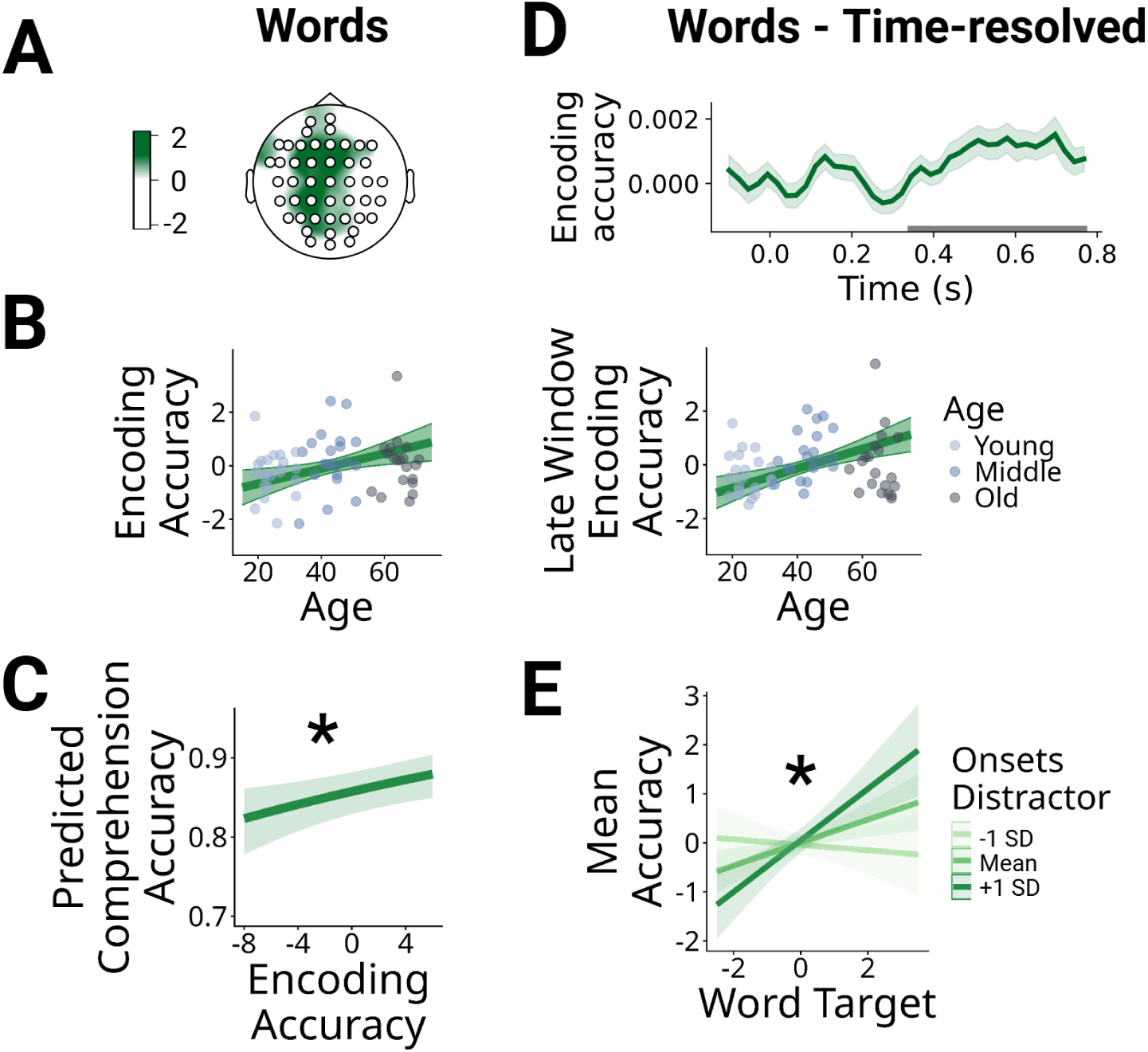
Neural tracking and neuro-behavioral correlations for the higher-level target features. A: Topographies for the encoding accuracies. Significant electrodes in mass-univariate t-tests are marked by white dots. B: Age effects on encoding accuracy. Significant effects are marked by regression lines. There was a positive correlation between age and word-level encoding accuracy. C: Neuro-behavioral correlations. Linear mixed model predictions of the effects of trial-wise neural encoding accuracy on comprehension accuracy. Positive effects of word-level target tracking on comprehension accuracy. D: Time-resolved encoding accuracy for the target stream and age effect on encoding accuracy in a late window (400 - 800 ms). Grey bars indicate significant encoding performance. E: Word-level tracking may compensate for increased distraction. Significant interaction between the tracking of word-level target features and distractor onsets (*Estimate* = 0.29*, SE* = 0.09*, t* = 3.09*, p_F DRcorrected_* = .008**). The positive effect of word-level target tracking on comprehension accuracy was larger at high levels of distractor onsets tracking, indicating a potential compensatory function of higher-level tracking.

#### Stronger higher-level tracking with increasing age

As displayed in Figure 3 B, there was a positive effect of age on higher-level target tracking at the word level (Estimate = 0.48, SE = 0.18, t = 2.69, p = .009**). Older adults therefore seem to display stronger neural responses to higher-level linguistic features.

#### Higher-level target tracking is correlated with improved comprehension

Consistent with previous results, the linear mixed model revealed positive effects of higher-level target tracking on comprehension performance. Thus, when word-level tracking of the target stream improved, comprehension performance was improved. The model predictions are visualized in Figure 3 C. There were no age interactions with this effect, indicating that this positive relationship was stable across the adult life span.

#### Higher-level linguistic tracking may compensate for increased distraction

In an explorative analysis, we tested the hypothesis that higher-level word tracking may compensate for increased neural tracking of the distractor stream. We used a multiple regression model predicting each subjects’ mean accuracy from the mean encoding accuracy of the word-level target features and the tracking of the distractor onsets and acoustics and their interactions. We additionally included three-way interactions between age, word-level target encoding accuracy, and lower-level distractor encoding accuracy. The analysis was controlled for the subject-specific SRT, the PTA, and the accuracy in the control condition. Results revealed a significant interaction between the word-level target and the onsets distractor encoding accuracies (*Estimate* = 0.29*, SE* = 0.09*, t* = 3.09*, p_F DRcorrected_*= .008**). This interaction indicates that the positive effect of word-level target tracking on comprehension performance was stronger when the distractor onset tracking was increased (see Figure 3 E). The effect was specific for the neural tracking of the distractor onsets, with no significant interaction emerging for distractor acoustic tracking. Thus, there seems to be a relationship between enhanced distractibility and higher-level tracking. This means that stronger higher-level tracking might compensate for increased individual distractibility. However, based on the available correlational evidence, no causal conclusions on the origin of this relationship can be drawn. There were no significant interactions between these effects and the participants’ age, which indicates that this compensatory mechanism may be a general strategy to maintain speech processing under challenging conditions across the adult life span. The full model outputs can be retrieved from Table S1.

#### Time-resolved encoding accuracies reveal late tracking for word-level features

To further investigate the time course of word-level feature encoding, we conducted an explorative analysis using time-resolved encoding accuracies for the word-level model variant. This was done to confirm that the observed effects result from late linguistic responses. Results revealed that the higher-level word features were represented in late time windows starting at 400 ms after word onset. This effect was specific to the target sentence. No significant encoding performance emerged for the distractor sentence. The results are visualized in Figure 3 D.

#### Stronger late linguistic tracking with increasing age

The results revealed a positive correlation between age and the encoding accuracy of word level target features in a late window starting at 400 ms after target word onset (Estimate = 0.62, SE = 0.16, t = 3.77, p < .001**). These late time windows have been commonly associated with higher-level processes related to word identity and the processing of word expectations. Thus, the neural tracking of higher-level processes seems to increase across the adult life span, suggesting a stronger reliance on higher-level processing with increasing age.

## Discussion

By combining detailed behavioral comprehension responses and multivariate, trial-resolved neural encoding models, we investigated age-related changes in the behavioral and neural mechanisms underlying speech comprehension in multitalker environments. The results revealed two main findings: on the one hand, older adults showed signs of impaired stream segregation abilities and enhanced susceptibility to distraction. This behaviorally relevant impairment likely contributes to age-related comprehension difficulties in multitalker environments. On the other hand, we found stronger behavioral and neural engagement with higher-level linguistic information with increasing age. Connecting these findings, we revealed evidence that higher-level tracking may compensate for stronger neural distractibility by facilitating successful comprehension.

Collectively, our results reveal a complex pattern of age-related changes in lower-level mechanisms and increased higher-level recruitment. Increased higher-level recruitment may serve a compensatory function for an individuals’ increased susceptibility to distraction. These results thus provide detailed insights into age-related impairments and potential compensatory strategies which help to maintain efficient speech processing across the adult life span.

### Comprehension is influenced by lower- and higher-level features

Behavioral results revealed that comprehension performance was influenced by lower-level word audibility, as well as word surprisal and word entropy. In addition, there was an interaction between word audibility and word surprisal indicating that the effect of surprisal was stronger at lower levels of audibility. The magnitude of this behavioral effect remained constant across the adult life span. Thus, on a word-by-word level, older adults were equally able to use acoustic and linguistic information in order to support speech comprehension.

Additionally, there was a positive effect of word entropy on comprehension performance, indicating that comprehension improved when uncertainty was higher. This effect got stronger with increasing age, which might indicate that dynamic modulation of attention in relation to uncertainty increases with age. This means that the allocation of attention in response to highly uncertain contexts, as well as the deallocation of attention in contexts deemed rather certain seems to be stronger in older adults. Thus, the context certainty seems to be a stronger predictor of older adults’ comprehension performance. In line with this conclusion, older adults seemed to benefit more from overall sentence context, when comparing comprehension performance in sentences and word lists. These results concur with behavioral effects suggesting a stronger reliance on contextual cues in older adults^13,24,25^.

### Neural tracking of lower- and higher-level information is positively correlated with comprehension

We investigated neural tracking for the target and distractor streams across several levels of the speech hierarchy, from phonetic and phonological (lower-level) mechanisms to higher-level probabilistic features. Neuro-behavioral correlations revealed that the trial-by-trial neural tracking of target word and phoneme onsets was positively associated with comprehension performance, consistently with previous results^47–52^. Additionally, we observed a positive effect of the neural tracking of distractor acoustics on comprehension. This effect suggests that an initial lower-level analysis of the distractor stream is beneficial for stream segregation. For a detailed discussion of this effect, see^46^.

Increased tracking of higher-level word features for the target stream was correlated with better comprehension performance. This is consistent with previous results^32,48,53–55^ and indicates that, as expected, when participants understood the sentences, this was reflected in a stronger neural tracking of the corresponding word-level features.

Our results did not reveal any interactions between age and the effects of acoustic or word-level encoding on comprehension, indicating that the effects of the encoding strength on comprehension performance remain constant across the adult life span. The reliance on acoustic and higher-level linguistic cues to parse the speech signal as the basis for speech comprehension thus seems to be stable across the adult life span.

### Increased distraction with increasing age

Using neuro-behavioral correlations on a trial-by-trial level, we showed that the neural tracking of distractor word and phoneme onsets had a negative effect on behavior in older adults but not in younger adults. Thus, poorer speech comprehension in older adults could be explicitly associated with impaired attentional filtering and decreased neural inhibition of the distractor stream^22^. Similar findings have been observed for hearing impaired populations^23,56,57^, suggesting that reduced stream segregation could be related to peripheral declines. Importantly, unlike previous studies^22,38,56^, we provide a missing link between the neural tracking of the distractor stream and the behavioral comprehension performance. By employing neuro-behavioral correlations on a trial-by-trial level, we show that the enhanced tracking of distractor onsets is detrimental to comprehension particularly in late adulthood. This effect emerged while task difficulty was individually adjusted on the basis of each individuals’ speech reception threshold. Thus, differences in the neural distractibility cannot be attributed to an overall poorer performance in older adults due to peripheral hearing constraints. This result further reinforces evidence indicating that impaired speech comprehension in older adults might rely on impaired attentional filtering^22,24,58^. This could be related to age-related declines in the ability to inhibit the neural processing of task-irrelevant information^22,38^.

### Increased higher-level recruitment with increasing age

A positive correlation between age and the strength of word-level encoding for the target stream indicates that the neural tracking of higher-level probabilistic features was stronger in older adults than in younger adults. Using a time-resolved measure of encoding accuracy, we found that, when word-level probabilistic features could reliably predict brain activity (in a late processing window starting at 400 ms after word onset), a positive association between age and word-level encoding performance emerged. This late processing window is typically associated with the processing of expectations and word identity^30,48,59–61^. Combined with our behavioral findings of an enhanced reliance on word entropy and sentence context to support speech comprehension, the results indicate an increased reliance on higher-level linguistic processing in late adulthood across the behavioral and neural levels. Consistent with previous behavioral and neural results^13,24,35^, these findings indicate that older adults might compensate for lower-level peripheral declines by using higher-level linguistic processing. This is supported by our evidence showing that increased word-level tracking can compensate for detrimental effects of a stronger distractor representation across the adult life span. Thus, an increased reliance on higher-level processing may serve as a strategy to maintain successful speech comprehension in challenging listening conditions across the adult life span^62,63^.

### Conclusion

In sum, we provide evidence for a behaviorally relevant decline in attentional filtering across the adult life span, which might underly speech comprehension difficulties in late adulthood. This lower-level impairment may be compensated for by an increased reliance on higher-level mechanisms at the behavioral and neural levels. Thus, older adults may increasingly rely on contextual cues provided in natural language to maintain speech comprehension abilities in late adulthood. These results provide implications for interventions targeting speech comprehension and social integration in late adulthood, arguing for enhancing higher-level cognitive processing to support compensational recruitment in speech comprehension.

## Materials and Methods

### Participants

Participants aged 19 to 71 years were recruited from the participant database maintained at the Max Planck Institute for Human Cognitive and Brain Sciences, Leipzig, Germany. Parts of the current sample were used in^46^. Participants received monetary compensation of 12 *€* per hour for their participation. The study was approved by the ethics committee of the medical faculty at Leipzig University (ethics vote: 032/24-ek). Inclusion criteria included age-appropriate psychosocial functioning as well as normal hearing. Psychosocial functioning was confirmed by a score of 27 or higher on the mini mental state examination^64^. For normal hearing, the criterion was a pure tone threshold below or equal to 25 dB HL across frequencies from 250 to 8000 Hz in at least one ear. 4 additional participants were excluded after screening due to insufficient hearing performance. The mean pure tone threshold for the remaining participants’ better ear in our sample was M = 8.12 dB HL (range = −4 - 25 dB HL). The full audiometric results for the included participants can be retrieved from Figure S1. All participants were German native speakers, right handed, and reported being neurologically and psychiatrically healthy. The final sample comprised 63 participants (35 women) aged 19 to 71 years (M = 43.75, SD = 17.15).

### Procedure

Participants listened to a total of 240 trials separated into 12 blocks comprised of 20 sentences each. Each trial consisted of two sentences presented in parallel. One sentence was spoken by a female speaker and the other one was spoken by a male speaker. Participants were instructed to follow the male speaker while ignoring the female speaker. After listening to the sentences and a brief period of 1 second to avoid EEG artifacts, participants were instructed to repeat the target sentence. In case they did not understand the complete sentence, they were instructed to repeat the part of the sentence that they understood. If they did not understand anything, they were instructed to say so. The behavioral paradigm is schematically shown in Figure 1. The experimental procedure is described in full detail in^46^. Experimental control was administered by the Psychophysics Toolbox Version 3 running on Matlab on a Windows machine^65,66^. The experiment was conducted in a sound-proof, electromagnetically shielded booth. Stimuli were presented using Koss KSC75 headphones clipped to the participants’ ears. Speech recordings were administered using a high quality condenser microphone (Rode NT55) at 44100 Hz. Sentences were presented at six individually adjusted target-to-distractor ratios [TDR; −2, −1, 0, 1, 2, 10 dB relative to their 50% speech reception threshold]. The last condition served as a control condition, in which the target sentence should be well comprehended by every participant and thus serving as a control for attention and working memory. Control trials were excluded from the neural analyses. The mean accuracy in the control trials was satisfyingly high in all participants (M = 97%, range = [91%, 99.5%]). The mean accuracy in the control trials was considered as a control variable for the neural analyses and the neuro-behavioral correlations. Participants additionally listened to one block of word lists composed of unrelated nouns presented at 0 dB TDR relative to their SRTs.

To control behavioral performance for peripheral differences in hearing abilities, speech reception thresholds (SRTs) were measured using an adaptive staircase procedure adapted from^67,68^ including 20 additional stimuli. Distractor intensity was held constant at 70 dB SPL. Target intensity was controlled by the staircase procedure, with a decrease in intensity if participants correctly repeated the full target sentence, and an increase in intensity if participants incorrectly repeated at least one word. Accuracy was rated by the experimenter. Similar as in^67^, the staircase procedure started with an TDR of 5 dB and a step size of 6 dB. Step size was decreased by a factor of 0.85 following each turning point. SRTs were calculated as the average of the TDR ratios preceding the final 5 turning points. The mean SRT across participants was *M* = −7.70(*SD* = 1.36).

### Stimuli

Stimuli were recorded at 44100 Hz on a high quality condenser microphone (Rode NT55) in a sound-proof booth. The distractor sentences started 0.5 seconds prior and lasted for a minimum of 0.3 seconds longer than the target sentences to ensure a complete masking of the target stream. Accordingly, the target sentences were comprised by 5-7 words and lasted for 2-4 seconds. The distractor sentences were comprised by 7-10 words and lasted for 3-5 seconds. The sentence material was retrieved from a German sentence corpus^69^ after filtering for the suitable sentence length and excluding sentences that contained names or very unusual words. Additionally, similar suitable sentences were created by GPT-3.5. The prompts used to create the sentences can be retrieved from Table S2. All sentences were loudness normalized to −23 dB LUFS using pyloudnorm^70^.

### EEG data acquisition

EEG was recorded in a soundproof, electromagnetically shielded booth at a sampling rate of 1000 Hz using a REFA8 68-channel amplifier system (TMSi, Oldenzaal, the Netherlands), grounded to the sternum. The recording included 63 Ag/AgCl electrodes positioned according to the 10-20 layout in ANT Neuro waveguard original caps as well as two mastoid external mastoid references (A1, A2). Eye movements were recorded using bipolar electrooculogram (EOG) electrodes placed at the outer sides of both eyes and at the top and bottom of the right eye. The EEG signals were monitored throughout the experiment.

### EEG preprocessing

EEG preprocessing was conducted using MNE python^71^ and followed recommendations for temporal response function (TRF) analyses^72^. The signal was filtered between 0.5 and 15 Hz and rereferenced to the average of the mastoid references. Bad channels were rejected manually based on a larger variance than neighboring channels. Independent component analysis (ICA) with 15 components was conducted to reduce artifacts from blinking and eye movements. The rejected components were selected automatically based on their correlation with the EOG electrodes. We finally interpolated bad channels using spherical splines^73^ and downsampled the signals to 128 Hz for TRF estimation to reduce computation times. The signal was epoched between target sentence onset and offset.

### Features

#### Acoustic features

Acoustic features included stimulus envelopes and envelope derivatives. Additionally, we investigated the neural tracking of word and phoneme onsets. All features were generated separately for target and distractor streams.

Envelopes and envelope derivatives were derived from auditory spectrograms calculated using naplib^74^. Spectrogram calculation involved a cochlear filter bank of 128 logarithmically-spaced constant-Q filters and a hair cell model to approximate the human peripheral auditory system. Broadband envelopes were calculated using the mean across all spectrogram channels. Envelope derivatives were derived using the half-wave rectified derivative of the broadband envelope^31^.

To derive word and phoneme onsets, stimuli and text transcripts were automatically aligned using forced alignment available in the WebMAUS Basic module of the BAS Web Services^75,76^. To avoid artifacts of target sentence onset, the first word onset in each sentence was included as a separate predictor and the remaining word-level predictors only included word onsets starting from the second word.

#### Word level features

Word frequency was derived from the SUBTLEX-DE subtitles corpus^77^. Word frequency was defined as the logarithm of word occurence per million, as provided in the SUBTLEX-DE database. Word surprisal is defined as the inverse probability of each word given the preceding sentence context^78,79^. Surprisal of word *i* is defined as the negative logarithm of word probability:

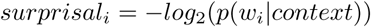

with (*p*(*w_i_*|*context*) being the word probability given the preceding sentence context derived from german-gpt2^80^. This model is an autoregressive causal language model based on the GPT-2 architecture, trained on large-scale German text corpora. The model relies on a GPT-2 byte-pair encoding (BPE) tokenizer. This means that words may be split into multiple subword tokens. Word entropy is defined as the uncertainty of the upcoming word given the previous sentence context from the previous word *w_i−_*_1_ to the first word in the sentence *w_i−n_*.

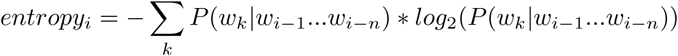

Word entropy values were derived from the probability distribution in German GPT-2^80^. Word-level surprisal and entropy were computed as the sum of token surprisals and entropies. All word-level predictors excluded the first word in each target sentence to avoid artifacts of sentence onset.

#### Phoneme level features

Phoneme surprisal reflects probability of each phoneme given the preceding phonemes in the current word. It is based on a probability prior captured by word frequency. Phoneme surprisal was defined as the inverse conditional probability of each phoneme, given the preceding phonemes in the same word:

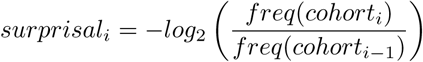

*Cohort_i_*is the cohort of possible words at the current phoneme and *freq*(*cohort*) is the sum of the word frequencies of all words in the cohort.

Phoneme entropy reflects the uncertainty about the next phoneme. Entropy at phoneme *i* is defined by:

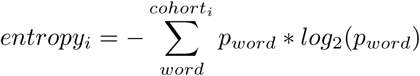

with *p_word_* being the probability of each word as reflected by its frequency. To appropriately separate the word and phoneme levels, we only used phoneme events that did not correspond to word onsets, i.e., omitting the first phonemes in each word. All features were min-max normalized to retain the natural zero baseline for the sparse regressors before entering the TRF.

#### Speaker characteristics

As mentioned above, target and distractor sentences were presented by a male and a female voice, respectively. There was only one male and one female speaker across the full experiment. To avoid introducing an age-related confound due to the stronger severity of age-related hearing loss in higher frequencies, we refrained from randomizing the assignment of the male and female speakers to the target and distractor roles. This was done to prevent confounding effects of the speakers’ pitch hight, potentially dampening the effects. To control for the differences introduced by the speakers, the analysis was controlled for fundamental frequency. Fundamental frequency was calculated using the pYIN algorithm^81,82^ available in the python package librosa^83^.

Additionally, we controlled the higher-level analyses for a set of higher level speaker characteristics. To obtain the speaker characteristics, a subsample of the participants (N = 35) took part in an online experiment after completing the EEG experiment. Here, participants were presented with ten sentences spoken by the same female and the male speakers used in the EEG experiment. After each sentence, they were asked to rate the speakers on a set of 7 characteristics: age, educatedness, dominance, attractiveness, health, and professionalism^84^. The characteristics were rated on 8 point ratings scales and the ratings were summarized across trials. The mean ratings were added into the acoustic model containing the respective characteristic at each word onset and the predicted responses were residualized for the estimation of the higher-level model.

#### TRF Models

To investigate the neural encoding of acoustic and linguistic features, we used multivariate TRFs. The analysis was implemented in mTRFpy^85^ using multiple linear regression with ridge regularization. The TRF is a kernel that describes the linear transformation from an ongoing stimulus to an ongoing neural response for a specified set of time lags^72^. When applying forward modeling, the TRF is used to predict the EEG signal from a set of stimulus features. To obtain the predicted EEG signal, the TRF is convolved with the speech features. The prediction accuracy is then obtained by correlating the predicted EEG signal with the actual signal. The prediction accuracy can then be interpreted as the amount of neural representation of the speech features. Feature representations were investigated in time lags from −100 to 800 ms relative to event onset.

TRF models were trained within participants. We used a 90-10 cross-validation with 50 randomly sampled folds following recent recommendations for evaluating brain decoding models^86^. For each fold, 20 trials were randomly selected to comprise the test set. Data from these trials was held out for the model training. We obtained trial-wise model fits for each individual trial from the TRF models trained across trials. Each trial appeared in the test set for 5 times. The mean of all 5 cross-validated model fits was used for all subsequent analyses. Model fit was defined as Pearson correlation between predicted and observed EEG across the full trial, averaged across sensors. The results did not depend on the exact choice of the cross-validation procedure (see Figures S3 and S4 for a reproduction of the results using a 10-fold cross-validation with independent folds).

Naturally, higher-level features are highly correlated to lower-level features in speech, making it important to control analyses investigating higher-level features for lower-level variance^34,87^. To control the higher-level analyses for lower-level responses, we residualized the lower-level responses from the EEG data used in the hlhigher-level analyses. This was important to isolate the unique contributions of the higher-level features, given the high collinearity between the lower-level and the higher-level features in natural speech. For the residualization, we conducted a separate TRF analysis predicting the EEG signals from all acoustic variables with no cross-validation. Model predictions from this analysis were then subtracted from the original EEG data to generate residualized EEG responses that do not contain responses resulting from acoustic features. All analyses involving higher-level probabilistic features were conducted on these residualized EEG responses. The regularization parameter was fitted separately for lower-level and higher-level models^88^. Fitting individual regularization parameters per participant led to of overfitting in the acoustic analyses and we therefore opted to fit the regularization parameters across participants. The optimal regularization parameter was determined based on the median across participants, as the mean regularization was strongly driven by a small number of outliers displaying very high regularization parameters. There was no correlation between age and the individual regularization parameters, suggesting that the model fitting procedure did not bias the age effects.

#### Statistical Inference

We used a shuffling approach to infer statistical significance of the TRF model fits. We grouped the individual features in order to avoid multicollinearity issues in regression model fitting. The model variants and their corresponding features are shown in Table 2. We conducted 599 shuffling iterations for each model variant following recommendations for iteration-based statistical inference^89^. In each iteration, we systematically shuffled the features of interest while keeping all other input features and model parameters identical. We used different shuffling methods for lower-level and higher-level models, corresponding to the characteristics of the respective features. For the acoustic models, trial labels were shuffled. For the higher-level models, we employed a more conservative approach of shuffling the values for each feature while preserving the onset times.

**Table 2:** Feature groups.

| Model variant | Shuffled features |
| --- | --- |
| Acoustic | Envelope, envelope derivative |
| Onsets | Word onsets, phoneme onsets |
| Phoneme level | Phoneme surprisal, phoneme entropy |
| Word level | Word surprisal, word frequency, word entropy |

Statistical significance was inferred by subtracting the mean shuffled model fits from the unshuffled model fits. Statistical inference was performed using mass-univariate one-tailed one sample t-tests with threshold-free cluster enhancement with 9999 permutations and a cluster forming threshold of *α* = 0.05. Adjacency was computed using Delaunay triangulation of sensor positions and the analysis was controlled for multiple comparisons separately for each feature and stream. Only the features displaying significant encoding performance in at least one sensor were used in the subsequent analyses. No significant tracking for the distractor higher-level features could be observed, indicating that these features do not show a generalizable effect on encoding performance. Thus, the distractor higher-level features were not used in any of the subsequent analyses, consistently with previous results on similar datasets^31^. In a previous study on the same data^46^, higher-level tracking for the distractor stream only emerged when isolating the incorrect trials.

#### Time-resolved encoding accuracy

To further disentangle the time scale of the word-level effects, we derived a time-resolved measure of encoding accuracy. In this analysis, the TRF was splitted into windows of 6 samples in steps of 3 samples. The TRF weights were set to zero outside of these windows, so that only certain sections of the TRF weights were used for the prediction of the EEG signal^90,91^.

#### Age effects on encoding accuracy

To investigate if the neural tracking of the investigated features increases with increasing age, we conducted multiple linear regression analyses predicting the subject-wise model fits from age. The analysis was controlled for the SRT and the mean accuracy in the main experiment and in the control condition by adding these variables as additional predictors. Due to the high correlation between age and hearing ability, as measured by the pure tone average (PTA), we added the orthogonalized PTA into the regression analyses. This means that only the share of variance in the PTA that could not be explained by age was added into the model. The correlations between age and the control variables are displayed in Figure S2.

#### Behavioral Analysis

The speech recordings were automatically transcribed offline using Whisper^92^. Subsequently, the transcriptions were manually controlled. We then assessed the overlap between the transcribed response and the target sentence, marking each word as accurately repeated or inaccurate.

To investigate the influence of lower-level and higher-level features on comprehension, word-by-word repetition performance was predicted from word surprisal, word entropy, and word audibility. The amount of masking varies on a moment-by-moment basis in competing speech stimuli, leading to some speech segments being relatively well audible. This has been described as glimpsing^93^. Glimpsed segments can inform the comprehension of segments that coincide with higher amounts of acoustic masking. To estimate the amount of available glimpses in each target word, we calculated the glimpse rate for each target word segmentation^93^. The glimpse rate relies on a comparison between the spectrogram amplitudes of the target and distractor streams. At each target word segmentation, we defined the proportion of spectrotemporal events that exceed a local target-to-distractor threshold. Based on previous work, this local threshold was set to −5 dB^93^.

To estimate the effects of word surprisal, word entropy, audibility, and their interaction on word repetition performance in the experimental conditions, we calculated a generalized linear mixed model with a logistic link function using lme4 in R^94^. The analysis was controlled for word frequency, word length, sentence length (in words), TDR condition, SRT, orthogonalized PTA, and control accuracy by adding these variables as additional predictors. We included random intercepts for subjects and sentences. To investigate the relationship between TRF model fits and behavioral performance, trial-wise model fits for all model variants were entered into the logistic regression model. We included interactions between age and all variables of interest (word surprisal, entropy, and audibility, TRF model fits). All variables were z-scored before entering the regression model. With regards to age, the original age values are used only for illustration purposes.

To rule out issues of multicollinearity, we calculated variance inflation factors using the R package car^95^. The variance inflation factors ranged from 1.00 to 3.80 and we therefore assume no issues of multicollinearity^96^. The highest variance inflation factors were observed for word surprisal and word frequency. However, despite their substantial correlation (r = -.67), we assume that there was sufficient non-redundant information in both predictor variables. The full model explained 20.6 % of variance in comprehension considering fixed effects only, and 50.3 % of variance if random effects were taken into account.

To quantify each individual’s gain from sentence context, mean performance in the main experiment was compared with the mean performance in an additional block of word lists. The words contained in the lists were selected from all nouns appearing in the sDeWac corpus^97^ at least 1000 times, with a length between 5 and 11 letters. We furthermore wanted to avoid involuntary similarities between nouns in the word lists, as this could have created a sense of coherence while listening to the lists of words. Therefore we chose the nouns that on average had the lowest similarities to all other candidate nouns. As a measure of semantic similarity we used cosine similarity among the corresponding word vectors in fastText, a widely used word embedding model^98^. To investigate age effects, we predicted the standardized difference scores from age, controlling for SRT, orthogonalized PTA, control accuracy, and mean accuracy in the main experiment using multiple linear regression.

#### Compensational effects of higher-level tracking

In an explorative analysis, we tested if word-level tracking can compensate for increased susceptibility for distraction. Therefore, we calculated the mean accuracy for each participant. This accuracy was predicted from the mean encoding accuracy for word-level target features and for distractor onsets and acoustics. We included three-way interactions between age, word-level target tracking, and distractor onsets and acoustic tracking. The analysis was controlled for the SRT, orthogonalized PTA and the accuracy in the control condition, serving as a proxy for working memory performance. P-values were corrected for multiple comparisons using FDR.

## Data availability

The data will be deposited to a suitable repository upon acceptance of the manuscript.

## Code availability

All custom code underlying the conclusions can be retrieved from https://github.com/vivienbarchet/compensage.

## Competing interests

The authors declare no competing interest.

## Author Contributions

AVB, AB, and GH designed the experiment; AVB oversaw data collection; AVB preprocessed and analyzed the data under supervision by AB, JMR, GH, and JO; AVB wrote the first draft; all authors contributed to the final version of the manuscript.

## Acknowledgments

We thank Heike Boethel and Jasmin Wend for their support in data acquisition and our participants for volunteering to participate in the study.

## Funding

AVB was supported by a PhD fellowship from the International Max Planck Research School (IMPRS) on Cognitive Neuroimaging. GH was supported by the Lise Meitner Excellence Program of the Max Planck Society, the German Research Foundation (DFG, HA 6314/4-2; Research Unit 5429/1 (467143400), HA 6314/10-2), and the European Research Council (ERC-2021-COG101043747). JO was supported by the German Research Foundation (DFG, OB 352/2-2). JMR was supported by the Max Planck Institute for Empirical Aesthetics.

## Notes

### Competing Interest Statement

The authors have declared no competing interest.

### Summary of Updates

Modified analyses, small changes in the methods and results sections.

