## supplementary material for "Linguistic information compensates for age-related decline in attentional filtering"

Alice Vivien Barchet [1,2,3,*], Andrea Bruera [1], Johanna M. Rimmele [4], Jonas Obleser [5,6], and Gesa Hartwigsen [1,2]

[1] Research Group Cognition and Plasticity, Max Planck Institute for Human Cognitive and Brain Sciences, Leipzig, Germany

[2] Cognitive and Biological Psychology, Leipzig University, Leipzig, Germany

[3] International Max Planck Research School on Cognitive NeuroImaging (IMPRS CoNI), Leipzig, Germany

[4] Department of Cognitive Neuropsychology, Max Planck Institute for Empirical Aesthetics, Frankfurt, Germany

[5] Department of Psychology, University of Lübeck, Lübeck, Germany

[6] Center of Brain, Behavior, and Metabolism, University of Lübeck, Lübeck, Germany

**Figure S1: Pure Tone Thresholds**

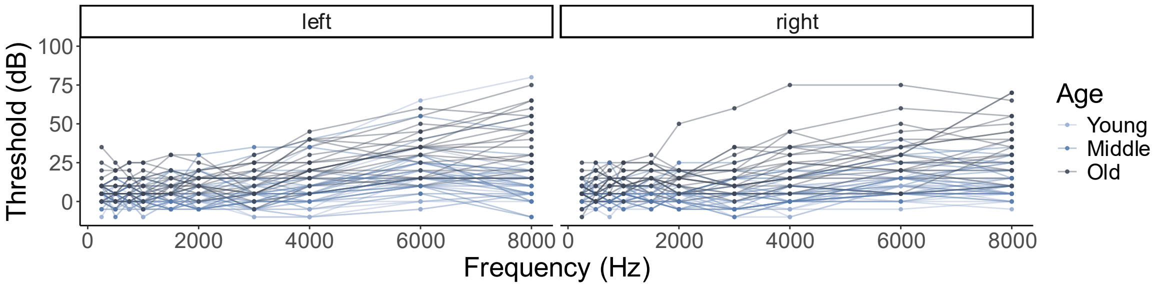


**Figure S2: Correlations between Age and Confounding Variables**

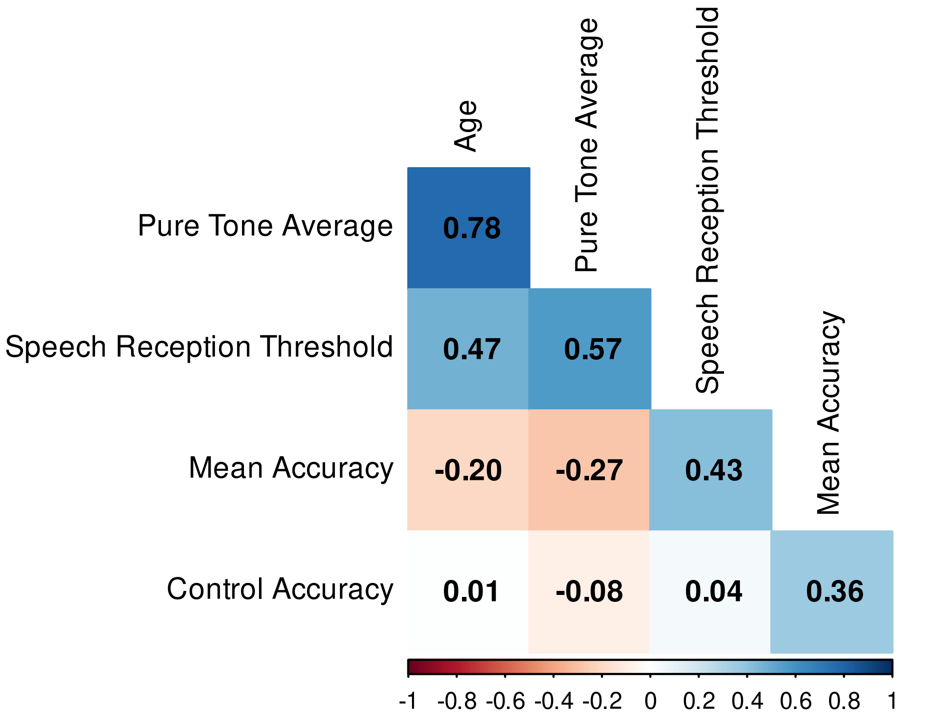

**Table S1: Model Outputs for the Compensation Analysis**

| | Estimate | Std.Error | CI | t | p |
|---|---|---|---|---|---|
| Age | -0.66 | 0.09 | [-0.84, -0.48] | -7.28 | <0.001** |
| Word Target | 0.23 | 0.08 | [0.07, 0.39] | 2.94 | .011* |
| Onsets Distractor | 0.05 | 0.09 | [-0.13, 0.22] | 0.51 | .707 |
| Acoustic Distractor | 0.07 | 0.09 | [-0.11, 0.25] | 0.75 | .624 |
| Word Target*Onsets Distractor | 0.29 | 0.09 | [0.1, 0.48] | 3.09 | .008** |
| Word Target*Acoustic Distractor | -0.18 | 0.12 | [-0.43, 0.06] | -1.48 | .224 |
| Age*Word Target | -0.01 | 0.09 | [-0.2, 0.17] | -0.13 | .901 |
| Age*Onsets Distractor | -0.17 | 0.08 | [-0.34, 0] | -2.01 | .093 |
| Age*Acoustic Distractor | 0.14 | 0.09 | [-0.05, 0.32] | 1.46 | .224 |
| Age*Word Target*Onsets Distractor | -0.04 | 0.1 | [-0.24, 0.17] | -0.37 | .763 |
| Age*Word Target*Acoustic Distractor | 0.07 | 0.12 | [-0.17, 0.32] | 0.61 | .681 |
| Control Accuracy | 0.27 | 0.08 | [0.11, 0.43] | 3.36 | .005** |
| PTA (residualized) | -0.41 | 0.09 | [-0.59, -0.23] | -4.62 | <0.001** |
| SRT | 0.3 | 0.04 | [0.23, 0.37] | 8.36 | <0.001** |

** $p < .01$, * $p < .05$ (FDR corrected)

**Table S2: GPT Prompts for target sentence generation**

| **Prompts** |
|---|
| Give me 10 sentences with 5-7 words that could stem from a story. |
| Generate 10 everyday sentences with 5-7 words. Only use frequent words and do not use commas. |
| Variate the sentence structure. Do not use question marks or commas. |
| Generate more complex sentences with 5-7 words. Only use frequent words and do not use commas. Variate the sentence structure. Do not use question marks or commas. |

Note. The same prompts were used for the distractor sentences, replacing 5-7 words with 7-10 words.

**Figure S3: Neural tracking and neuro-behavioral correlations for the lower-level features resulting from a 10-fold cross-validation**

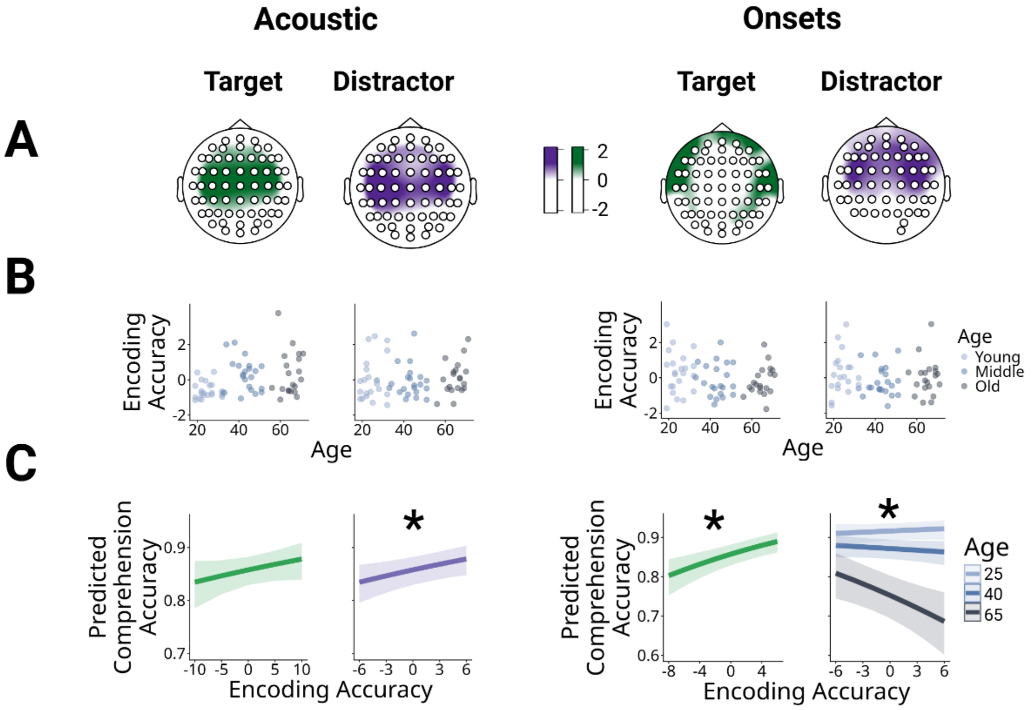


**Figure S4: Neural tracking and neuro-behavioral correlations for the higher-level target features resulting from a 10-fold cross-validation**

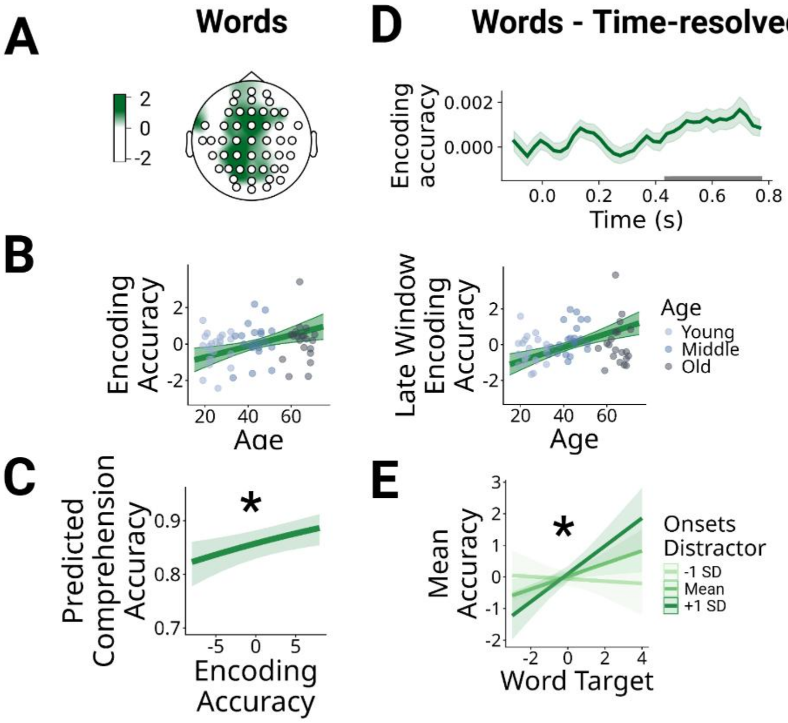
